# Cryo-ET reveals nanoscale thick filament disorganization in MYH7 P710R hypertrophic cardiomyopathy cardiomyocytes

**DOI:** 10.64898/2026.02.18.706593

**Authors:** Magda Zaoralová, Joseph Yoniles, Prerna Giri, Richard G. Held, Alison Vander Roest, Peter D. Dahlberg, Daniel Bernstein, Alexander R. Dunn, Leeya Engel

## Abstract

Hypertrophic cardiomyopathy (HCM) is the most common monogenic inherited heart disease and is a major cause of sudden death in individuals under 35 years of age. HCM is associated with progressive tissue-level disarray and subcellular disorganization in individual cardiomyocytes. Mutations in β-cardiac myosin (MYH7), the second most common genetic cause of HCM, commonly result in changes in sarcomeric force production, but how this leads to altered cell- and tissue-level organization is unclear. Here, we use cryo-electron tomography (cryo-ET) to bridge the molecular and cellular scales by visualizing the nanoscale organization of individual myosin-containing thick filaments within sarcomeres of human induced pluripotent stem cell-derived cardiomyocytes (hiPSC-CMs). Compared to isogenic wild-type controls, hiPSC-CMs expressing P710R MYH7 exhibit pronounced disruption of the hexagonal packing of thick filaments within individual sarcomeres, which would be difficult to visualize using conventional light microscopy or even room-temperature electron microscopy. We also observe ribosome infiltration into areas of sarcomeric disorder for both wild-type and P710R MYH7 hiPSC-CMs, suggesting that disordered regions may be sites of local proteostasis or remodeling. Together, these data illuminate how altered myosin activity can propagate to yield dramatic changes in sarcomeric organization in HCM.

## Introduction

Hypertrophic cardiomyopathy (HCM) affects 1 in 500 individuals and is linked to an increased risk of heart failure and sudden cardiac death (1). Physiologically, HCM is characterized by hypercontractility and hypertrophy (thickening) of the left ventricular wall, diastolic dysfunction, and myocardial interstitial fibrosis. Over 1,000 genetic mutations are linked to HCM, with the second most commonly mutated gene, *MYH7*, encoding the β-cardiac myosin heavy chain (2–6). Mutations in MYH7 result in increased force production at the level of individual cardiomyocytes (CMs) (7). However, the mechanisms that link changes in force production to pathological hypertrophy remain incompletely understood (8, 9).

Force production in CMs depends on the contraction of myofibrils, which are made up of repeating structural units termed sarcomeres. Each sarcomere consists of myosin-containing thick filaments and thin filaments templated by filamentous (F)-actin. The barbed, (+) ends of actin filaments are anchored at the Z-discs, massive protein assemblies that link neighboring sarcomeres. The processes by which sarcomeres form termed sarcomerogenesis, are not fully understood (10). Although the underlying molecular mechanisms are debated, recent studies using human induced pluripotent stem cell (hiPSC)-CMs as a model system reveal that sarcomeres develop from actin- and myosin-containing stress fibers, and that this process depends on myosin-generated force (11, 12). How and whether sarcomerogenesis may be altered in the context of HCM is, to our knowledge, little explored.

In addition to hypertrophy, HCM is accompanied by CM disarray, in which the normally orderly, parallel organization of sarcomeres is disrupted. This disarray may be clinically significant, as it may result in conduction irregularities that lead to ventricular arrhythmias and sudden death (13). While the mechanistic origins are not well understood, emerging evidence suggests that CM disarray may precede hypertrophy (14). CM-level disarray is accompanied by increased myofibril disorder and an increase in the width of the Z-discs as assessed by conventional electron microscopy (EM) (15). This increase in Z-disc width is recapitulated in hiPSC-CMs gene edited to express mutations in MYH7 that cause HCM (16). These cells also demonstrate hypertrophy relative to isogenic, control hiPSC-CMs, increased force production at the cellular level, and altered Ca^2+^ handling, illustrating that they recapitulate key cellular features of HCM.

The P710R mutation in MYH7 leads to severe, pediatric onset HCM (17). This mutation is associated with an increase in force production in CMs even after correction for increased CM volume due to cellular hypertrophy (16). How P710R and other myosin mutations alter cell and tissue-level organization is not yet fully understood. At the molecular level, increased force production is thought to arise from a decrease in the proportion of myosin heads that occupy an inactivated conformation termed the super-relaxed state (SRX), which increases the number of myosin heads available for force production per sarcomeric length (18, 19). Previous studies revealed that the P710R MYH7 mutation leads to disorganization at the level of cardiomyocytes and myofibrils (20), and that myofibril-level disorganization is recapitulated in P710R MYH7 hiPSC-CMs (16). Importantly, these and other studies were performed using conventional electron microscopy on resin-embedded cardiomyocytes. Although this modality provides insight into cellular and subcellular organization, it lacks molecular resolution and involves preparation techniques that can introduce distortions (*i*.*e*., fixation, resin embedding). Conversely, biochemical studies have provided detailed mechanistic insight into how mutations in MYH7 alter myosin conformation and catalytic activity, but lack cellular context (6, 19).

Here we pair cryo-electron tomography (cryo-ET), leveraging grid micropatterning techniques, with recent advances in the culture and maturation of hiPSC-CMs to investigate how P710R MYH7, and by extension other HCM-associated mutations in β-cardiac myosin, alter sarcomere organization at the nanometer scale. Cryo-ET provides an unparalleled ability to visualize nanoscale structures such as the cytoskeleton in a near-native environment (21–30) and has recently been employed to reveal sarcomere ultrastructure in isolated mouse muscle myofibrils (31–33) and to reconstruct sub-nanometer resolution structures of the human thin filament in a cellular context (34). Here we find that P710R mutation results in disorganization in the thick filaments that propagates to the sarcomere, myofibril, and cellular scales. In addition, we observe infiltration of ribosomes in the disordered regions of both HCM and matched control hiPSC-CMs. These data provide insight into how molecular-scale perturbations resulting in HCM propagate to changes on the cellular length scale.

## Results

We developed a specialized micropatterning workflow to prepare the hiPSC-CMs for cryogenic focused ion beam milling (cryo-FIB) and cryo-ET (34) (Fig. 1A). This workflow promoted the attachment of hiPSC-CMs to specific locations on the surface of the grid to facilitate cryo-FIB thinning and cryo-ET imaging of the cells. Positioning cells in the centers of the grid squares also improved the freezing efficiency by preventing cell clustering, which generally led to the formation of crystalline ice during the freezing process. Together, these improvements considerably increased the number of areas accessible for cryo-FIB sample preparation. In addition, the use of rectangular adhesion patterns induced the hiPSC-CMs to adopt an elongated shape that replicates the shape of CMs *in vivo* and promotes CM maturation (35).

**Fig. 1.**
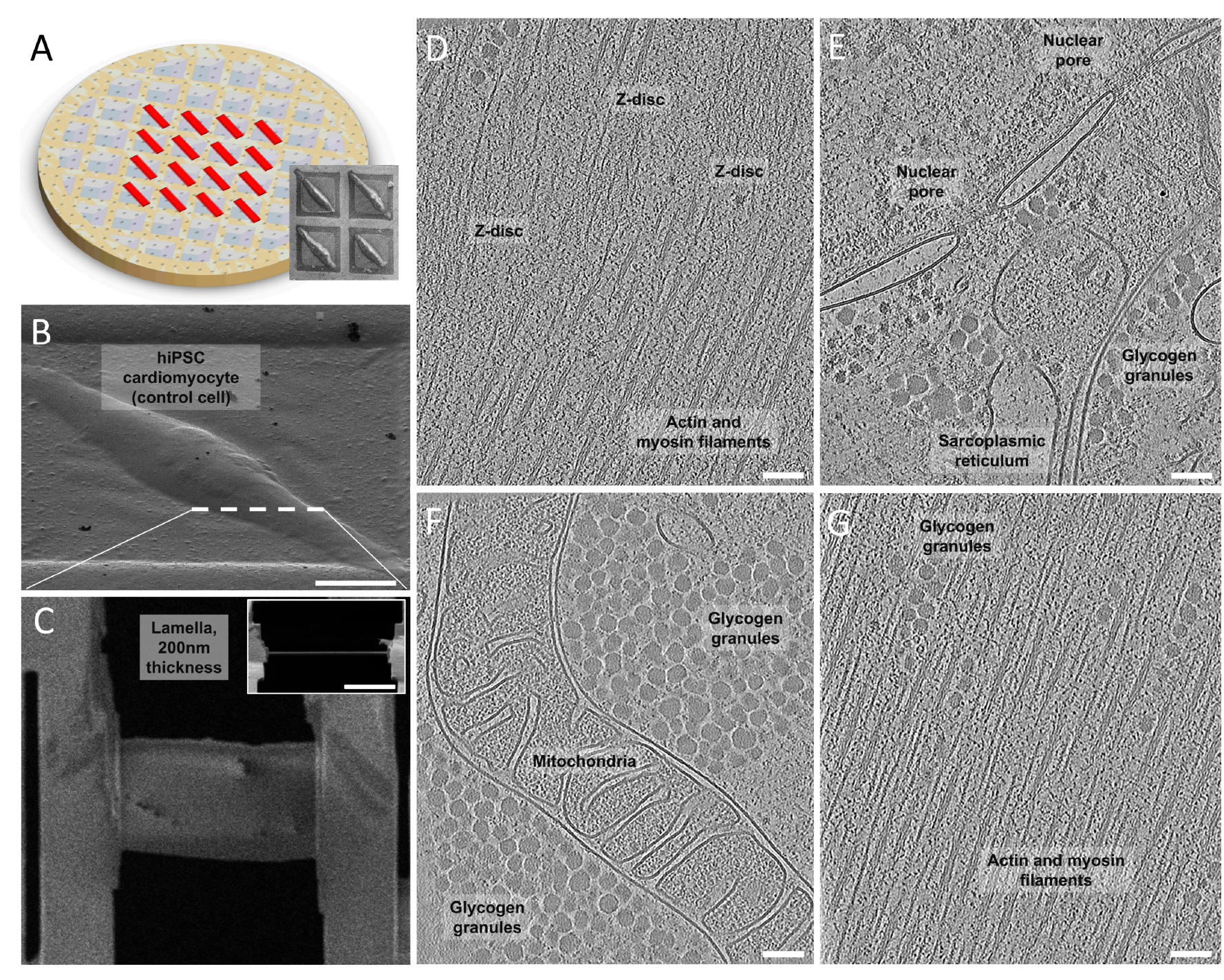
Overview of workflow for cryo-FIB and cryo-ET of hiPSC-CMs. (A) Schematic of micropatterned electron microscopy grids (Matrigel micropatterns in red). Insert shows cryo-SEM of vitrified patterned hiPSC-CMs. Scale bar, 100 um. (B) Well positioned and shaped cardiomyocyte. Red dashed line shows the rought location of lamella targetting. Scale bar, 20 um. (C) An example of a lamella prepared by cryo-FIB and imaged via SEM with an approximate thickness of 200 nm. Insert shows cross section of lamella imaged by FIB, scale bar, 5 um. (D) Slices of the reconstructed 3D volume of the lamellae, showing an array of cellular features: Z-discs, nuclear pore, glycogen granules, sarcoplasmic reticulum, mitochondria, and thick and thin filaments. Scale bars, 100 nm.

Using FIB milling, vitrified cells were thinned to approximately 200 nm. The positions of the lamellae were chosen so that regions adjacent to the nucleus were selected (Fig. 1B). The lamellae were imaged at lower magnification (12k-fold magnification) to capture the overview of the section and provide context for high, 52k-fold magnification tomographic imaging of selected regions. Following TEM data collection, denoised tomographic reconstructions were computed and segmented for features of interest. This workflow was applied to both P710R MYH7 and isogenic controls, allowing for the comparison of subcellular and molecular-scale structures preserved in their native, hydrated state.

We observed striking differences in sarcomere organization between the P710R and control cells, quantified by the angular distribution of the myosin thick filaments within the sarcomeres (Fig. 2). The P710R mutant hiPSC line showed a significantly greater average dispersion angle than the control cells, where dispersion angle is defined as the angle between each thick filament and the average orientation of thick filaments in the tomogram (p=5.7×10^*−*10^). This sub-sarcomeric *myofilaments* disarray is distinct from the classic disarray of *myofibrils* that has been observed in TEM images of fixed and resin-embedded heart tissue, which manifests at length scales an order of magnitude larger than is characterized in this study. To our knowledge, sub-sarcomeric disorder in orientation of the individual thick-filaments in the context of HCM constitutes a novel observation that could not have been straightforward to achieve using conventional electron microscopy techniques.

**Fig. 2.**
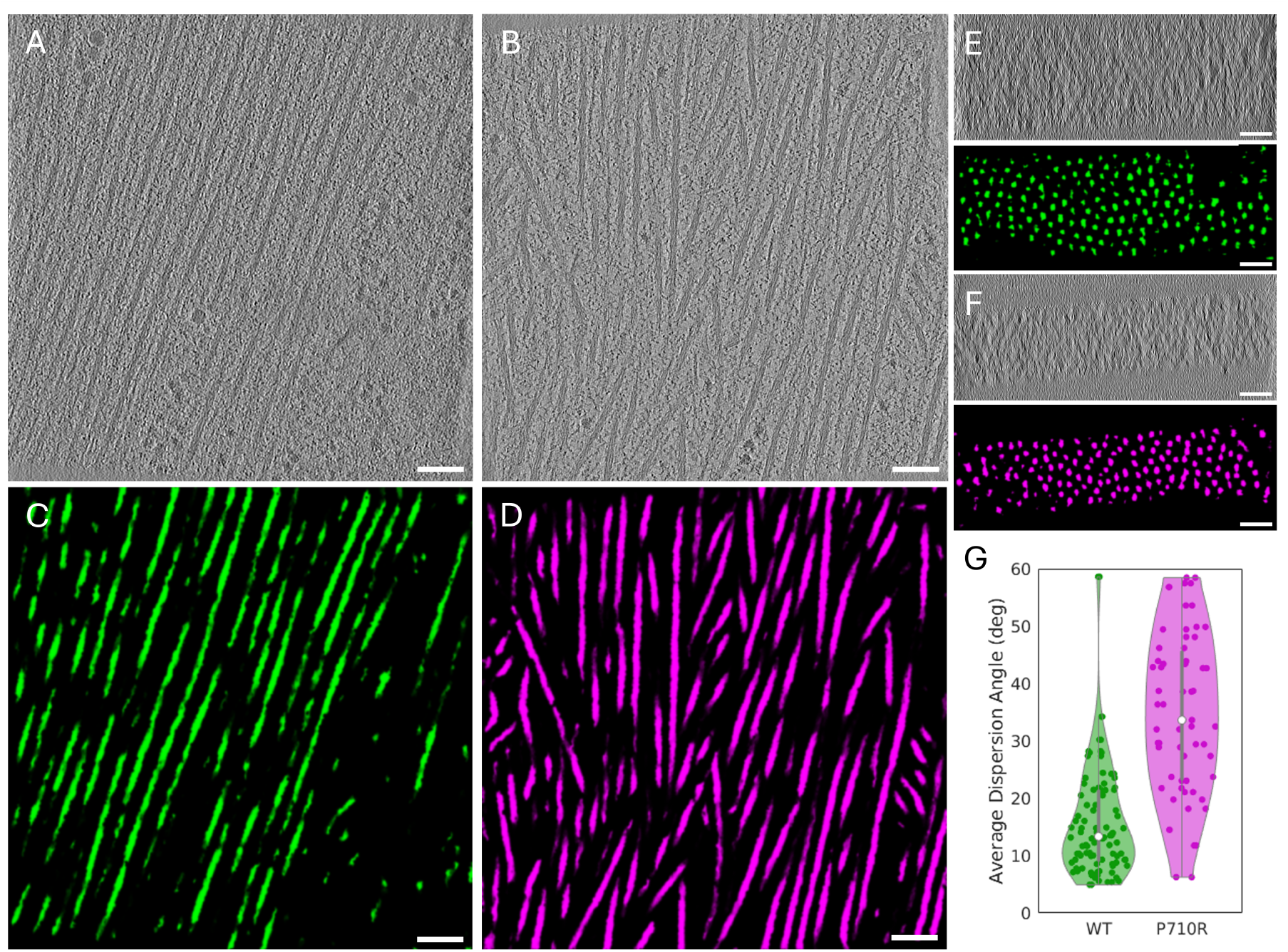
Tomographic slices of control (A) and P710R (B) hiPSC-CMs. Segmentation of the thick filaments in the control (C) and mutant (D) cells are visualized in green and magenta, respectively. (E,F) Cross sections of the tomograms for the control (E) and mutant (F) cells with the segmented thick filaments visualized in the panels below. (G) Average dispersion angle, quantifying the filament disorder. P-value calculated using student’s t test comparing grid replicates: p=5.7×10^*−*10^ . (See Supplementary Movie for full tomogram). Scale bars, 100 nm.

In addition to thick and thin filaments, we observed a higher number of ribosomes localized within disorganized sarcomeric regions in P710R MYH7 hiPSC-CMs as compared to control cells (Fig. 3, see Discussion). Our cryo-tomography data also included a wide array of cellular compartments and features in both the P710R MYH7 hiPSC-CMs and corresponding control cells. Among these were Z-discs, ribosomes, nuclear pores, glycogen granules, sarco- or endoplasmic reticulum, and mitochondria (Fig. 1). The general sub-cellular organization observed in the cryo-lamellae was comparable in both cell lines. Interestingly, we observed a high degree of variability in Z-disc shape in both the mutant and control cell lines (see Fig. 1D for tomographic slice of Z-disc in control cells). We did not distinguish any obvious differences in mitochondria or other organelles between the two cell lines. However, due to the involvement of mitochondria in generating energy for the cell, these organelles could be a promising target for future studies using our approach.

**Fig. 3.**
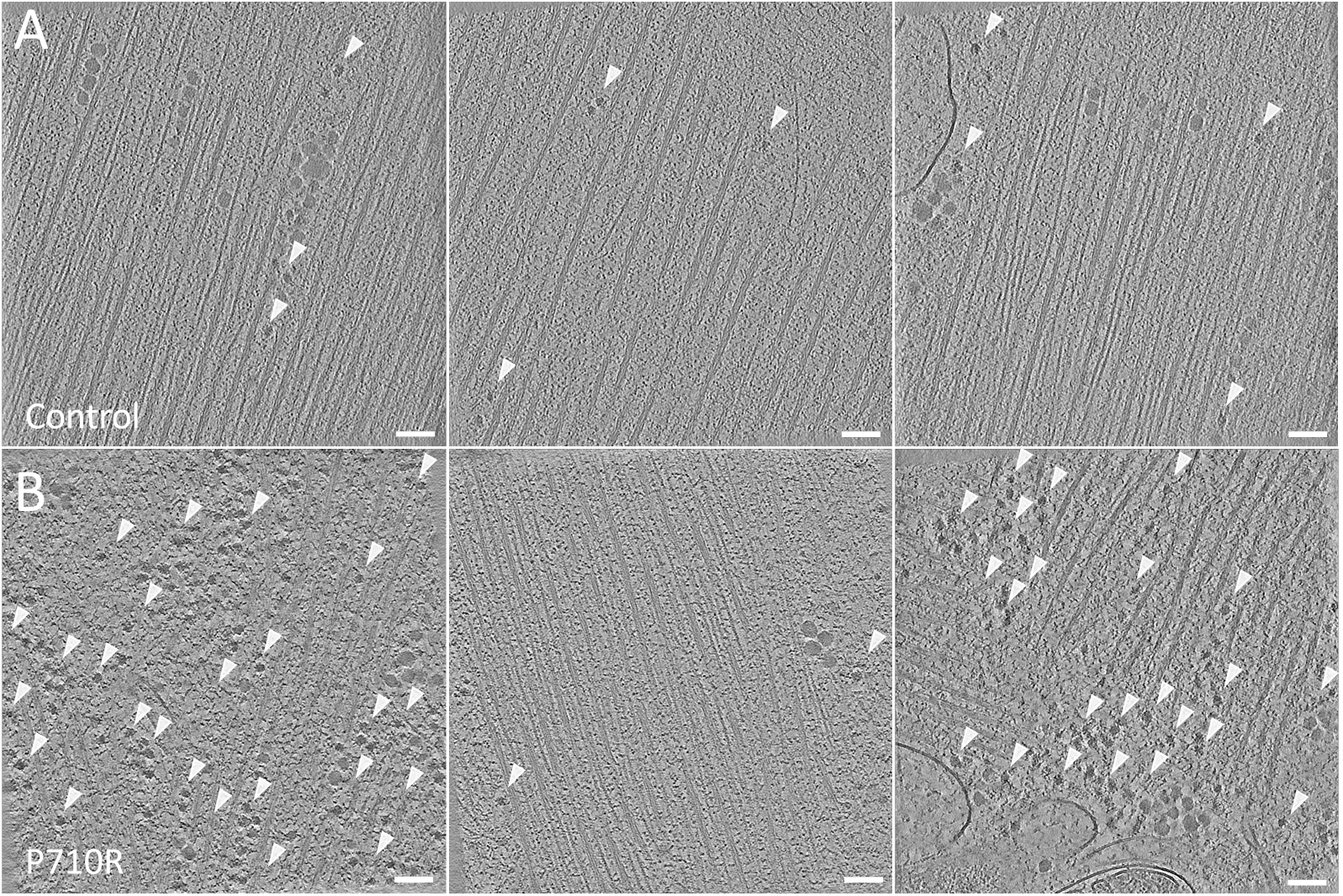
Ribosome distribution and abundance within sarcomeres. (A) Control cells with organized sarcomeric regions show a low prevalence of ribosomes, while (B) P710R mutant cell sections show a higher degree of disorganization of the sarcomeres and striking up-regulation of ribosomes in disordered regions. Scale bars for all tomographic slices, 100 nm.

## Discussion

While the genetic basis of HCM is well established, the intermediate mechanisms linking molecular perturbations to cellular and tissue-level pathology remain incompletely understood. Despite operating through distinct molecular mechanisms, HCM mutations in sarcomeric proteins, converge on a common outcome of hypercontractility (36–41). A prevailing hypothesis suggests that increased force production triggers mechanosensitive signaling cascades that lead to thickening of cardiac muscle and impaired cardiac function. P710R MYH7 cells demonstrate this with significantly elevated force production, and inhibition of downstream ERK and Akt signaling is sufficient to prevent the hypertrophic phenotype (16). Clinical support for a causal role for hyper-contractility comes from evidence that removal of the thickened heart wall by myectomy or alcohol septal ablation not only relieves symptoms, but also significantly reduces further thickening and future complications (42, 43). Supporting the hypothesis of a shared propagation mechanism across a diverse set of mutations, the small-molecule drug mavacamten, which works by decreasing the ATPase activity of myosin heads, increases the SRX ratio while decreasing sarcomeric force production without targeting any specific mutation location (44). Our cryo-ET data address this gap in understanding by examining the critical ultrastructural changes that may precede, or drive, overt hypertrophy.

The hiPSC-CMs most closely resemble fetal CMs, a developmental stage that precedes overt clinical manifestations of HCM by years to decades. In this context, our observation that the P710R MYH7 mutation increases disorder in the orientation of individual thick filaments relative to matched controls suggests that mutation-driven changes in the contractile apparatus may emerge early during cardiomyocyte maturation, well before hypertrophy becomes apparent. We speculate that nanoscale disruption of thick-filament organization may precede and potentially drive larger-scale sarcomeric disarray and hypertrophy that are the hallmarks of HCM. Defining how altered myosin activity gives rise to the nanoscale disorder we describe will be an important focus for future work.

The increased local concentration of ribosomes in disorganized sarcomeric regions complements an earlier study reporting that mRNA encoding for MYH7 localized to the sarcomeres, with MYH7 mRNA appearing to qualitatively correlate with less organized regions of the sarcomeres. We speculate that the ribosomes we observe may facilitate local translation of myosin, and possibly other proteins at the sarcomere. If so, such a model would be consistent with studies showing that both the production and proteolytic turnover of sarcomeric proteins are tightly regulated. In one such example, it was reported that the inhibition of the ubiquitin ligase led to an abundance of free-floating myosin, resulting in the disruption of the sarcomeric assembly and a disarray of myofibrils (45–47). Our finding complements these and other studies by providing precise locations of ribosomes and confirms their increased concentration and varied distribution within disordered sarcomeres. In concert with the aforementioned studies, our data suggest that the increased ribosome concentration within disordered regions of the sarcomere and local translation of myosin could lead to or be caused by disarray within the sarcomere.

While our cryo-ET approach provides valuable insights into sub-sarcomeric organization in the context of HCM, several considerations should be noted. The hiPSC-CM model used here, may not capture all features relevant to the disease, which occurs in the context of a multicellular tissue with complex physiology. Despite recent advances in hiPSC-CM maturation showing mature isoforms of MYH7, the internal cellular organization remains immature compared to adult heart cardiomyocytes, with certain features like transverse tubules undeveloped or absent. Additionally, our analysis captures only a single timepoint in sarcomere assembly and maturation, which represents a single snapshot of a dynamic process. Future cryo-ET studies examining sub-sarcomeric changes in HCM heart tissue across multiple developmental stages would complement these findings and provide deeper insights into disease progression. The workflow developed here can be readily applied to such investigations, as we have demonstrated the ability to visualize key proteins within the close-to-native, cryo-preserved cellular context.

## Methods

### EM grid preparation

Cells were seeded on UltrAufoil EM grids (200 mesh R2/2, Quantifoil Micro Tools, GmbH) (48) that were micropatterned with 17 × 118 µm rectangles to achieve a 1:7 aspect ratio, known to promote a more mature cardiac phenotype in hiPSC-CMs (35). Micropatterning was performed using Alveole PRIMO maskless photopatterning system following our earlier studies (49–52) (see Fig. 1A). Briefly, the EM grids were immobilized on silicone sheeting (Specialty Manufacturing Inc.) mounted on glass coverslips, exposed to atmospheric plasma, incubated in 0.01% Poly-L-Lysine (CAS Number 25988-63-0, p4707 Sigma) overnight, rinsed, and then incubated in a 100 mg/mL solution of mPEG-Succinimidyl Valerate (PEG-SVA, MW 5,000 Laysan Bio Inc) in 0.1 HEPES (pH 8.5) for 1 hr. The grids were then sterilized in 70% ethanol, rinsed, and incubated in a 1:150 dilution of Matrigel in DMEM/F12 for 1 hour at 37 degrees Celsius.

### Cell culture

The control hiPSC line (SCVI-113) was obtained from the Stanford Cardiovascular Institute iPSC Biobank. The mutant hiPSC line, P710R, was generated using CRISPR/Cas9 gene editing as described in our previous paper (16). Both the control and mutant (P710R) hiPSCs were differentiated into cardiomyocytes following Allen Institute standard operating procedures and with some modifications, as previously described (16). Briefly, hiPSCs were cultured on tissue culture plates coated with Matrigel (356231, Corning) and grown for 4 days in mTESR media (85850, Stemcell Technologies) before being differentiated into hiPSC-CMs. On day 52, control and mutant cells were lifted with TrypLE and seeded onto the micropatterned EM grids. First the Matrigel solution was aspirated from the grids and replaced with replating media (RPMI + B27 + 10 % Knock-out Serum Replacement + 10 µM Rock inhibitor). Next, a cell suspension containing approximately 500 cells was added to each grid. After two hours, additional media was added to each dish for a 24 hour incubation period after which media was replaced with RPMI + B27 and the cells were cultured for seven days on the EM grids prior to plunge freezing. The cells were vitrified in a Leica EM GP2 plunge freezer. The grids were then clipped into autogrid rings (Thermo Fisher Scientific) and stored in liquid nitrogen storage dewar prior to thinning by cryo-FIB.

### FIB-milling

Following plunge freezing, vitrified cells on EM grids were loaded into a cryo-FIB/SEM system Aquilos (Thermo Fisher Scientific) and coated with three layers of platinum. First, a rough sputter coating (at a default setting of 10 mA, 10 Pa and 15 s) using an integrated plasma coater. Second, approximately 500 nm of organoplatinum (GIS) was deposited, followed by second rough sputter coating. The cells were milled to approximately 180-210 nm thickness using a Gallium ion source accelerated by 30kV in the Aquilos system, starting with 1 - 0.5 nA for rough milling to 100 - 30 pA current for thinning and polishing. Maps and AutoTEM software (Thermo Fisher Scientific) were used for navigation, rough milling, and medium milling and polishing of selected regions. The level of automation and manual interventions varied depending on a lamella shape, size, and milling progress. During the final steps, the milling progress and final lamella shape were monitored by imaging with an electron beam of 2kV and 13pA. Polished lamellae were coated with fine sputter layer with 7 mA, 10 Pa and 7s coating conditions.

### TEM and tomographic data acquisition

Grids with prepared lamellae were cryo-transferred to a Thermo Fisher Scientific Krios G2 equipped with a Falcon 4i direct detector and Selectris X energy filter. Tilt series were collected using a Thermo Fisher Scientific Tomography 5 SW with integrated multi-shot tomogram acquisition, with a dose-symmetric tilt scheme, with tilt span ranging from 50-60 degrees and 2-3 degree tilt step increments. Region of interest was automatically tracked and autofocusing procedure was run for each tilt. An energy slit of 10 eV was used for the all data collection, including overview images of the lamella area. The dose was calibrated to 3.5 electron per Angstrom^2^ per projection, resulting in 140 e^*−*^/Å^2^ per tomogram. The pixel size of the final data acquisition was 2.2 and 2.3 Å and defocus values ranged from 2 to 5 µm. Data were acquired in the eer mode of the Falcon4i detector (53).

### Data processing

For data reconstruction, tilt series in eer format were converted to tiff file format using Relion3 implementation of SBGrid (ref). Individual projections were motion corrected with the MotionCor2, while simultaneously dividing the dataset into even/odd frames (28). Frames were aligned and reconstructed with AreTomo (27). Reconstructed 3D volume was used for model training in CryoCare (25, 26) and resulting model served as a denoising platform for the final 3D tomographic volume that was segmented and visualized using ORS Dragonfly neural networks and IMOD(29, 30).

## Supporting information

Tomogram_segmented_200nm_scale_bar

## CRediT author statement

**Magda Zaoralová**: Conceptualization, Methodology, Software, Formal analysis, Investigation, Data curation, Writing - Original Draft, Visualization, Project administration. **Joseph Yoniles**: Software, Formal analysis, Data curation, Writing - Review & Editing, Visualization. **Prerna Giri**: Investigation, Resources. **Richard G. Held**: Software, Formal analysis, Data curation, Writing - Review & Editing, Visualization. **Alison Vander Roest**: Conceptualization, Methodology, Investigation, Resources, Writing - Review & Editing. **Peter D. Dahlberg**: Resources, Writing - Review & Editing, Supervision, Funding acquisition. **Daniel Bernstein**: Resources, Writing - Review & Editing, Supervision, Funding acquisition. **Alexander R. Dunn**: Conceptualization, Writing - Original Draft, Supervision, Project administration, Funding acquisition. **Leeya Engel**: Conceptualization, Methodology, Investigation, Writing - Original Draft, Supervision, Project administration, Funding acquisition.

## ACKNOWLEDGEMENTS

This work was supported by NIH R35 GM130332 (ARD), NIH RM1 GM131981 and AHA 24CSA1257409 (DB), a Diane and Guilford Glazer Foundation Faculty Fellowship (LE), a Michigan-Israel Partnership Award (LE and AVR), the Arc Institute Fellowship (MZ), and the Panofsky Fellowship at the SLAC National Accelerator Laboratory as part of the Department of Energy Laboratory Directed Research and Development program under contract DE-AC02-76SF00515 (PDD). Cryo-FIB and cryo-ET data collection for this study were performed at the Stanford-SLAC Cryo-ET Specimen Preparation Center (SCSC), which is supported by the National Institutes of Health Common Fund’s Transformative High Resolution Cryoelectron Microscopy program (U24GM139166), and at the Stanford CryoEM Center (cEMc). The content is solely the responsibility of the authors and does not necessarily represent the official views of the NIH. This manuscript was prepared using the Henriques lab bioRxiv template on Overleaf. The authors declare no competing financial interests.

## Bibliography

1. Connie W. Tsao, Aaron W. Aday, Zaid I. Almarzooq, Alvaro Alonso, Andrea Z. Beaton, Marcio S. Bittencourt, Amelia K. Boehme, Alfred E. Buxton, April P. Carson, Yvonne Commodore-Mensah, Mitchell S.V. Elkind, Kelly R. Evenson, Chete Eze-Nliam, Jane F. Ferguson, Giuliano Generoso, Jennifer E. Ho, Rizwan Kalani, Sadiya S. Khan, Brett M. Kissela, Kristen L. Knutson, Deborah A. Levine, and et al. Lewis. Heart disease and stroke statistics-2022 update: A report from the american heart association. Circulation, 145(8): E153–E639, 2022.

2. Norbert Frey, Mark Luedde, and Hugo A. Katus. Mechanisms of disease: Hypertrophic cardiomyopathy. Nature Reviews Cardiology, 9(2):91–100, 2011. doi: 10.1038/nrcardio.2011.159.

3. Carolyn Y Ho, Sharlene M Day, Euan A Ashley, Michelle Michels, Alexandre C Pereira, Daniel Jacoby, Allison L Cirino, Jonathan C Fox, Neal K Lakdawala, James S Ware, et al. Genotype and lifetime burden of disease in hypertrophic cardiomyopathy: insights from the sarcomeric human cardiomyopathy registry (share). Circulation, 138(14):1387–1398, 2018.

4. Ali J Marian and Eugene Braunwald. Hypertrophic cardiomyopathy: genetics, pathogenesis, clinical manifestations, diagnosis, and therapy. Circulation research, 121(7):749–770, 2017.

5. J.G. Seidman and Christine Seidman. The genetic basis for cardiomyopathy: from mutation identification to mechanistic paradigms. Cell, 104(4):557–567, 2001. doi: 10.1016/s0092-8674(01)00242-2.

6. James A. Spudich. Three perspectives on the molecular basis of hypercontractility caused by hypertrophic cardiomyopathy mutations. Pflügers Archiv - European Journal of Physiology, 471(5):701–717, 2019. doi: 10.1007/s00424-019-02259-2.

7. Masataka Kawana, James A. Spudich, and Kathleen M. Ruppel. Hypertrophic cardiomyopathy: Mutations to mechanisms to therapies. Frontiers in Physiology, 13, 2022. doi: 10.3389/fphys.2022.975076.

8. Pamela A. Harvey and Leslie A. Leinwand. Cellular mechanisms of cardiomyopathy. Journal of Cell Biology, 194(3):355–365, 2011. doi: 10.1083/jcb.201101100.

9. Elizabeth A. Woodcock and Scot J. Matkovich. Cardiomyocytes structure, function and associated pathologies. The International Journal of Biochemistry amp;amp; Cell Biology, 37(9):1746–1751, 2005. doi: 10.1016/j.biocel.2005.04.011.

10. Elisabeth Ehler and Mathias Gautel. The sarcomere and sarcomerogenesis. Advances in Experimental Medicine and Biology, page 1–14, 2008. doi: 10.1007/978-0-387-84847-1_1.

11. Aidan M Fenix, Abigail C Neininger, Nilay Taneja, Karren Hyde, Mike R Visetsouk, Ryan J Garde, Baohong Liu, Benjamin R Nixon, Annabelle E Manalo, Jason R Becker, et al. Muscle-specific stress fibers give rise to sarcomeres in cardiomyocytes. Elife, 7:e42144, 2018.

12. Anant Chopra, Matthew L Kutys, Kehan Zhang, William J Polacheck, Calvin C Sheng, Rebeccah J Luu, Jeroen Eyckmans, J Travis Hinson, Jonathan G Seidman, Christine E Seidman, et al. Force generation via β-cardiac myosin, titin, and α-actinin drives cardiac sarcomere assembly from cell-matrix adhesions. Developmental cell, 44(1):87–96, 2018.

13. Gherardo Finocchiaro, Nabeel Sheikh, Ornella Leone, Joe Westaby, Francesco Mazzarotto, Antonis Pantazis, Cecilia Ferrantini, Leonardo Sacconi, Michael Papadakis, Sanjay Sharma, et al. Arrhythmogenic potential of myocardial disarray in hypertrophic cardiomyopathy: genetic basis, functional consequences and relation to sudden cardiac death. EP Europace, 23(7):985–995, 2021.

14. Patricia Garcia-Canadilla, Andrew C Cook, Timothy J Mohun, Onyedikachi Oji, Saskia Schlossarek, Lucie Carrier, William J McKenna, James C Moon, and Gabriella Captur. Myoarchitectural disarray of hypertrophic cardiomyopathy begins pre-birth. Journal of Anatomy, 235(5):962–976, 2019.

15. E. Bajusz, G. Rona, International Study Group for Research in Cardiac Metabolism, and South African Medical Research Council. Cardiomyopathies. Number v. 10 in Cardiomyopathies. University Park Press, 1973. ISBN 9783541061914.

16. Alison Schroer Vander Roest, Chao Liu, Makenna M Morck, Kristina Bezold Kooiker, Gwanghyun Jung, Dan Song, Aminah Dawood, Arnav Jhingran, Gaspard Pardon, Sara Ranjbarvaziri, et al. Hypertrophic cardiomyopathy β-cardiac myosin mutation (p710r) leads to hypercontractility by disrupting super relaxed state. Proceedings of the National Academy of Sciences, 118(24):e2025030118, 2021.

17. Juan Pablo Kaski, Petros Syrris, Maria Teresa Tome Esteban, Sharon Jenkins, Antonios Pantazis, John E Deanfield, William J McKenna, and Perry M Elliott. Prevalence of sarcomere protein gene mutations in preadolescent children with hypertrophic cardiomyopathy. Circulation: Cardiovascular Genetics, 2(5):436–441, 2009.

18. Melanie A. Stewart, Kathleen Franks-Skiba, Susan Chen, and Roger Cooke. Myosin atp turnover rate is a mechanism involved in thermogenesis in resting skeletal muscle fibers. Proceedings of the National Academy of Sciences, 107(1):430–435, 2010. doi: 10.1073/pnas.0909468107.

19. M. Kawana, S. S. Sarkar, S. Sutton, K. M. Ruppel, and J. A. Spudich. Biophysical properties of human β-cardiac myosin with converter-domain mutations that cause hypertrophic cardiomyopathy. Science Advances, 3:e1601959, 2017. doi: 10.1126/sciadv.1601959.

20. Sara Ranjbarvaziri, Kristina B Kooiker, Mathew Ellenberger, Giovanni Fajardo, Mingming Zhao, Alison Schroer Vander Roest, Rahel A Woldeyes, Tiffany T Koyano, Robyn Fong, Ning Ma, et al. Altered cardiac energetics and mitochondrial dysfunction in hypertrophic cardiomyopathy. Circulation, 144(21):1714–1731, 2021.

21. Ohad Medalia, Igor Weber, Achilleas S Frangakis, Daniela Nicastro, Günther Gerisch, and Wolfgang Baumeister. Macromolecular architecture in eukaryotic cells visualized by cryoelectron tomography. Science, 298(5596):1209–1213, 2002.

22. Shoh Asano, Benjamin D Engel, and Wolfgang Baumeister. In situ cryo-electron tomography: a post-reductionist approach to structural biology. Journal of molecular biology, 428 (2):332–343, 2016.

23. Lu Gan and Grant J Jensen. Electron tomography of cells. Quarterly reviews of biophysics, 45(1):27–56, 2012.

24. Saikat Chakraborty, Marion Jasnin, and Wolfgang Baumeister. Three-dimensional organization of the cytoskeleton: a cryo-electron tomography perspective. Protein Science, 2020.

25. Tim-Oliver Buchholz, Alexander Krull, Réza Shahidi, Gaia Pigino, Gáspár Jékely, and Florian Jug. Content-aware image restoration for electron microscopy. Methods in cell biology, 152:277–289, 2019.

26. Tim-Oliver Buchholz, Mareike Jordan, Gaia Pigino, and Florian Jug. Cryo-care: content-aware image restoration for cryo-transmission electron microscopy data. In 2019 IEEE 16th International Symposium on Biomedical Imaging (ISBI 2019), pages 502–506. IEEE, 2019.

27. Shawn Zheng, Georg Wolff, Garrett Greenan, Zhen Chen, Frank GA Faas, Montserrat Bárcena, Abraham J Koster, Yifan Cheng, and David A Agard. Aretomo: An integrated software package for automated marker-free, motion-corrected cryo-electron tomographic alignment and reconstruction. Journal of Structural Biology: X, 6:100068, 2022.

28. Shawn Q Zheng, Eugene Palovcak, Jean-Paul Armache, Kliment A Verba, Yifan Cheng, and David A Agard. Motioncor2: anisotropic correction of beam-induced motion for improved cryo-electron microscopy. Nature methods, 14(4):331–332, 2017.

29. James R Kremer, David N Mastronarde, and J Richard McIntosh. Computer visualization of three-dimensional image data using imod. Journal of structural biology, 116(1):71–76, 1996.

30. Jessica E Heebner, Carson Purnell, Ryan K Hylton, Mike Marsh, Michael A Grillo, and Matthew T Swulius. Deep learning-based segmentation of cryo-electron tomograms. Journal of Visualized Experiments (JoVE), (189):e64435, 2022.

31. Zhexin Wang, Michael Grange, Thorsten Wagner, Ay Lin Kho, Mathias Gautel, and Stefan Raunser. The molecular basis for sarcomere organization in vertebrate skeletal muscle. Cell, 184(8):2135–2150, 2021.

32. Zhexin Wang, Michael Grange, Sabrina Pospich, Thorsten Wagner, Ay Lin Kho, Mathias Gautel, and Stefan Raunser. Structures from intact myofibrils reveal mechanism of thin filament regulation through nebulin. Science, 375(6582):eabn1934, 2022.

33. Davide Tamborrini, Zhexin Wang, Thorsten Wagner, Sebastian Tacke, Markus Stabrin, Michael Grange, Ay Lin Kho, Martin Rees, Pauline Bennett, Mathias Gautel, and Stefan Raunser. Structure of the native myosin filament in the relaxed cardiac sarcomere. Nature, 623:863–871, 2023. doi: 10.1038/s41586-023-06690-5.

34. Rahel A Woldeyes, Masataka Nishiga, Alison S Vander Roest, Leeya Engel, Prerna Giri, Gabrielle C Montenegro, Andrew C Wu, Alexander R Dunn, James A Spudich, Daniel Bernstein, et al. Cryo-electron tomography reveals the structural diversity of cardiac proteins in their cellular context. bioRxiv, 2023.

35. Alexandre JS Ribeiro, Yen-Sin Ang, Ji-Dong Fu, Renee N Rivas, Tamer MA Mohamed, Gadryn C Higgs, Deepak Srivastava, and Beth L Pruitt. Contractility of single cardiomyocytes differentiated from pluripotent stem cells depends on physiological shape and substrate stiffness. Proceedings of the National Academy of Sciences, 112(41):12705–12710, 2015.

36. Masahiko Hoshijima. Mechanical stress-strain sensors embedded in cardiac cytoskeleton: Z disk, titin, and associated structures. American Journal of Physiology-Heart and Circulatory Physiology, 290(4):H1313–H1325, 2006.

37. Ruth F Sommese, Jongmin Sung, Suman Nag, Shirley Sutton, John C Deacon, Elizabeth Choe, Leslie A Leinwand, Kathleen Ruppel, and James A Spudich. Molecular consequences of the r453c hypertrophic cardiomyopathy mutation on human β-cardiac myosin motor function. Proceedings of the National Academy of Sciences, 110(31):12607–12612, 2013.

38. Arjun S Adhikari, Kristina B Kooiker, Saswata S Sarkar, Chao Liu, Daniel Bernstein, James A Spudich, and Kathleen M Ruppel. Early-onset hypertrophic cardiomyopathy mutations significantly increase the velocity, force, and actin-activated atpase activity of human β-cardiac myosin. Cell reports, 17(11):2857–2864, 2016.

39. Julien Robert-Paganin, Daniel Auguin, and Anne Houdusse. Hypertrophic cardiomyopathy disease results from disparate impairments of cardiac myosin function and auto-inhibition. Nature communications, 9(1):4019, 2018.

40. Robert C Cail, Bipasha Barua, Faviolla A Báez-Cruz, Donald A Winkelmann, Yale E Goldman, and E Michael Ostap. A myosin hypertrophic cardiomyopathy mutation disrupts the super-relaxed state and boosts contractility by enhanced actin attachment. Proceedings of the National Academy of Sciences, 122(52):e2521561122, 2025.

41. Neha Nandwani, Debanjan Bhowmik, Camille Glaser, Matthew Carter Childers, Rama Reddy Goluguri, Aminah Dawood, Michael Regnier, Anne Houdusse, James A Spudich, and Kathleen M Ruppel. Hypertrophic cardiomyopathy mutations y115h and e497d disrupt the folded-back state of human β-cardiac myosin allosterically. Nature Communications, 16(1):8751, 2025.

42. Nicholas G. Smedira, Bruce W. Lytle, Harry M. Lever, Jeevanantham Rajeswaran, Gita Krishnaswamy, Ryan K. Kaple, Diana O.W. Dolney, and Eugene H. Blackstone. Current effectiveness and risks of isolated septal myectomy for hypertrophic obstructive cardiomyopathy. The Annals of Thoracic Surgery, 85(1):127–133, 2008. ISSN 0003-4975. doi: 10.1016/j.athoracsur.2007.07.063.

43. Steve R. Ommen, Barry J. Maron, Iacopo Olivotto, Martin S. Maron, Franco Cecchi, Sandro Betocchi, Bernard J. Gersh, Michael J. Ackerman, Robert B. McCully, Joseph A. Dearani, Hartzell V. Schaff, Gordon K. Danielson, A. Jamil Tajik, and Rick A. Nishimura. Long-term effects of surgical septal myectomy on survival in patients with obstructive hypertrophic cardiomyopathy. Journal of the American College of Cardiology, 46(3):470–476, 2005. ISSN 0735-1097. doi: 10.1016/j.jacc.2005.02.090.

44. Robert L Anderson, Darshan V Trivedi, Saswata S Sarkar, Marcus Henze, Weikang Ma, Henry Gong, Christopher S Rogers, Joshua M Gorham, Fiona L Wong, Makenna M Morck, et al. Deciphering the super relaxed state of human β-cardiac myosin and the mode of action of mavacamten from myosin molecules to muscle fibers. Proceedings of the National Academy of Sciences, 115(35):E8143–E8152, 2018.

45. Yair E. Lewis, Anner Moskovitz, Michael Mutlak, Joerg Heineke, Lilac H. Caspi, and Izhak Kehat. Localization of transcripts, translation, and degradation for spatiotemporal sarcomere maintenance. Journal of Molecular and Cellular Cardiology, 116:16–28, 2018. ISSN 0022-2828. doi: 10.1016/j.yjmcc.2018.01.012.

46. Thomas G. Martin and Jonathan A. Kirk. Under construction: The dynamic assembly, maintenance, and degradation of the cardiac sarcomere. Journal of Molecular and Cellular Cardiology, 148:89–102, 2020. doi: 10.1016/j.yjmcc.2020.08.018.

47. Roua Hassoun, Heidi Budde, Saltanat Zhazykbayeva, Melissa Herwig, Marcel Sieme, Simin Delalat, Nusratul Mostafi, Kamilla Gömöri, Melina Tangos, Muhammad Jarkas, et al. Stress activated signalling impaired protein quality control pathways in human hypertrophic cardiomyopathy. International journal of cardiology, 344:160–169, 2021.

48. Christopher J Russo and Lori A Passmore. Ultrastable gold substrates for electron cryomicroscopy. Science, 346(6215):1377–1380, 2014.

49. Leeya Engel, Guido Gaietta, Liam P Dow, Mark F Swif, Gaspard Pardon, Niels Volkmann, William I Weis, Dorit Hanein, and Beth L Pruitt. Extracellular matrix micropatterning technology for whole cell cryogenic electron microscopy studies. Journal of Micromechanics and Microengineering, 29(11):115018, 2019.

50. Leeya Engel, Claudia G Vasquez, Elizabeth A Montabana, Belle M Sow, Marcin P Walkiewicz, William I Weis, and Alexander R Dunn. Lattice micropatterning for cryo-electron tomography studies of cell-cell contacts. Journal of structural biology, 213(4):107791, 2021.

51. Leeya Engel, Richard Held, Claudia G Vasquez, William I Weis, and Alexander R Dunn. Micropatterning em grids for cryo-electron tomography of cells. 2022.

52. Leeya Engel, Magda Zaoralová, Momei Zhou, Alexander R Dunn, and Stefan L Oliver. Extracellular filaments revealed by affinity capture cryogenic-electron tomography. Nature Communications, 16(1):9802, 2025.

53. Hui Guo, Erik Franken, Yuchen Deng, Samir Benlekbir, Garbi Lezcano, Bart Janssen, Lingbo Yu, Zev Ripstein, Yong Tan, and John Rubinstein. Electron-event representation data enable efficient cryoem file storage with full preservation of spatial and temporal resolution. IUCrJ, 7, 08 2020. doi: 10.1107/S205225252000929X.

